# Unsupervised learning and clustered connectivity enhance reinforcement learning in spiking neural networks

**DOI:** 10.1101/2020.03.17.995563

**Authors:** Philipp Weidel, Renato Duarte, Abigail Morrison

## Abstract

Reinforcement learning is a learning paradigm that can account for how organisms learn to adapt their behavior in complex environments with sparse rewards. However, implementations in spiking neuronal networks typically rely on input architectures involving place cells or receptive fields. This is problematic, as such approaches either scale badly as the environment grows in size or complexity, or presuppose knowledge on how the environment should be partitioned. Here, we propose a learning architecture that combines unsupervised learning on the input projections with clustered connectivity within the representation layer. This combination allows input features to be mapped to clusters; thus the network self-organizes to produce task-relevant activity patterns that can serve as the basis for reinforcement learning on the output projections. On the basis of the MNIST and Mountain Car tasks, we show that our proposed model performs better than either a comparable unclustered network or a clustered network with static input projections. We conclude that the combination of unsupervised learning and clustered connectivity provides a generic representational substrate suitable for further computation.

## 1 INTRODUCTION

Neural systems learn from past experience, gradually adapting their properties according to processing requirements. Be it in a merely sensory-driven situation or in high-level decision making processes, a key component of learning is to develop adequate and usable internal representations, allowing the system to assess, represent and use the current state of the environment in order to take actions that optimize expected future outcomes (Sutton & Barto, 2018).

While these principles have been extensively exploited in the domain of machine learning, leading to highly proficient information processing systems, the similarities with the biophysical reality are often merely conceptual. The standard approach to learning in neural systems has been end-to-end supervised training with error backpropagation (LeCun et al., 2015; Kriegeskorte & Golan, 2019). However, despite the proficiency of these algorithms, their plausibility under biological constraints is highly questionable (Nikolić, 2017; Marcus, 2018, but see e.g. Marblestone et al., 2016; Lillicrap & Santoro, 2019; Richards et al., 2019 for counterarguments). Whereas a growing body of literature has focused on bridging this divide by adjusting error backpropagation to make it more biophysically compatible (e.g. Sacramento et al., 2018; Bellec et al., 2019; Whittington & Bogacz, 2019), there remains a disconnect between these approaches and real neuronal and synaptic dynamics.

Biological neural networks operate with discrete pulses (spikes) and learn without explicit supervision. Synaptic efficacies are adjusted to task demands relying on local information, i.e. the activity of pre- and post-synaptic neurons, as well as unspecific, neuromodulatory gating factors (Porr & Wörgötter, 2007; Frémaux & Gerstner, 2016). As such, any model of the acquisition of internal representations ought to comply with these (minimal) criteria, gradually shaping the system’s properties to learn an adequate, task-driven partition of the state-space in a manner that allows the system to operate under different task constraints. Furthermore, these learning processes ought to ensure generalizability, allowing the same circuit to be re-used and operate on different input streams, extracting the relevant information from them and acquiring the relevant dynamical organization according to processing demands, in a self-organized manner.

Modelling studies have employed a variety of strategies to allow the system to internally represent the relevant input features (also referred to as environmental states, in the context of reinforcement learning). This can be done, for example, by manually selecting neuronal receptive fields (Potjans et al., 2009, 2011; Jitsev et al., 2012; Frémaux et al., 2013) according to a pre-specified partition of the environment or by spreading the receptive fields uniformly, in order to cover the entire input space (Frémaux et al., 2013; Jordan et al., 2017). These example solutions have major conceptual drawbacks. Manually partitioning the environmental state space is by definition an ad hoc solution for each task; whereas uniformly covering the whole input is a more generic solution, it can only be achieved for relatively low-dimensional input spaces. In both cases, the researcher imposes an assumption about the appropriate resolution of partitioning for a given task, and thereby implicitly also affects the learning performance of the neural agent. Both approaches are thus inflexible and restricted in their applicability.

It seems parsimonious to assume that the development of adequate internal states capturing relevant environmental features emerges from the way in which the input is projected onto the circuit. In the reservoir computing paradigm, the input projection acts as a non-linear temporal expansion (Schrauwen et al., 2007; Lukoševičius & Jaeger, 2009). Relatively low-dimensional input streams are thus non-linearly projected onto the circuit’s high-dimensional representational space. Through this expansive transformation, the neural substrate can develop suitable dynamic representations (Duarte & Morrison, 2014; Duarte et al., 2018) and resolve non-linearities such that classes that are not linearly separable in the input space can be separated in the system’s representational space. This property, commonly referred to as the *kernel trick*, relies on the characteristics of the neural substrate, acting as a non-linear operator, and the ensuing input-driven dynamics (Maass et al., 2002).

The output of reservoir computing models is commonly a simple (typically linear) supervised readout mechanism, trained on the circuit’s dynamics to find the pre-determined input-output mappings. Naturally, such supervised readouts are not intended to constitute realistic models of biological learning (Schrauwen et al., 2007), but instead constitute a metric to evaluate the system’s processing capabilities. In a biological system, one would expect the output projections of a reservoir to adapt in response to local information such as pre- and post-synaptic activity, possibly incorporating a global, diffusive neuromodulatory signal. A complication here is that synaptic learning that largely depends on the spiking activity of a pair of neurons is inevitably susceptible to the stochastic nature of that activity.

In this manuscript, we introduce a novel class of spiking neural network model that addresses the above issues with respect to partitioning the input space and learning output projections in a biologically plausible fashion. Our model consists of an input layer, a representation layer and and output layer. The representation layer is based on a balanced random network of spiking integrate-and-fire neurons including clusters as described in Rost et al. (2018); Rostami et al. (2020). When combined with unsupervised learning (Tetzlaff et al., 2013) of the input projections, the clusters become specialized, in a self-organized fashion, for features of the input space, thus allowing stable representations of the input to emerge that support a linear separation. The low-dimensional dynamics of the clustered network (Litwin-Kumar & Doiron, 2014) provide a stable basis for the three-factor, dopamine-modulated plasticity rule implemented by the output projections to learn appropriate input-output mappings using a reinforcement learning strategy.

We first demonstrate, using the XOR task, the capacity of the unsupervised learning rule to resolve non-linearities in the input, allowing the representation layer to generate linearly separable activity. We then investigate the performance of the full model on a 3-digit MNIST task. We show that unsupervised learning in the input projections allows the output projections to learn the correct classifications with a high degree of accuracy even though the learning is driven by a reinforcement (rather than supervisory) signal. The presence of the clusters in the representation layer cause the network to converge more quickly, and to a higher performance, than the corresponding unclustered network. The clustered network with plastic input projections resolves the non-linearities, captures the intra- and inter-class variance and elegantly deals with the highly overlapping inputs of this challenging task.

Finally, we test the model in a closed loop scenario on a task defined in continuous space and time with sparse rewards: the Mountain Car problem provided by the OpenAI Gym (Brockman et al., 2016). Once again, the clustered network with plastic input projections learns the task more effectively than an unclustered network or one with static input projections.

Notably, the network model is configured almost identically for these two quite dissimilar tasks, differing only in the mechanism by which the reinforcement learning signal is generated. In particular, the number of clusters is not optimized for the task, and the initial mapping of inputs to the representation layer is random. We thus conclude that the three components of our network model, namely, unsupervised learning on the input projections, clustered connectivity in the representation layer, and reinforcement learning on the output projection, combine to create a system possessing substantial generic learning capacity, requiring neither previous knowledge on how to partition the input space, nor training with biologically unrealistic supervised approaches. We anticipate that the components of unsupervised learning and clustered connectivity could also be employed to amplify the performance of other learning network models.

## 2 METHODS

### 2.1 Network Architecture

The network model consists of three layers, as illustrated schematically in Figure 1A. A complete tabular specification of the network and its parameters can be found in the supplementary materials.

**Figure 1.**
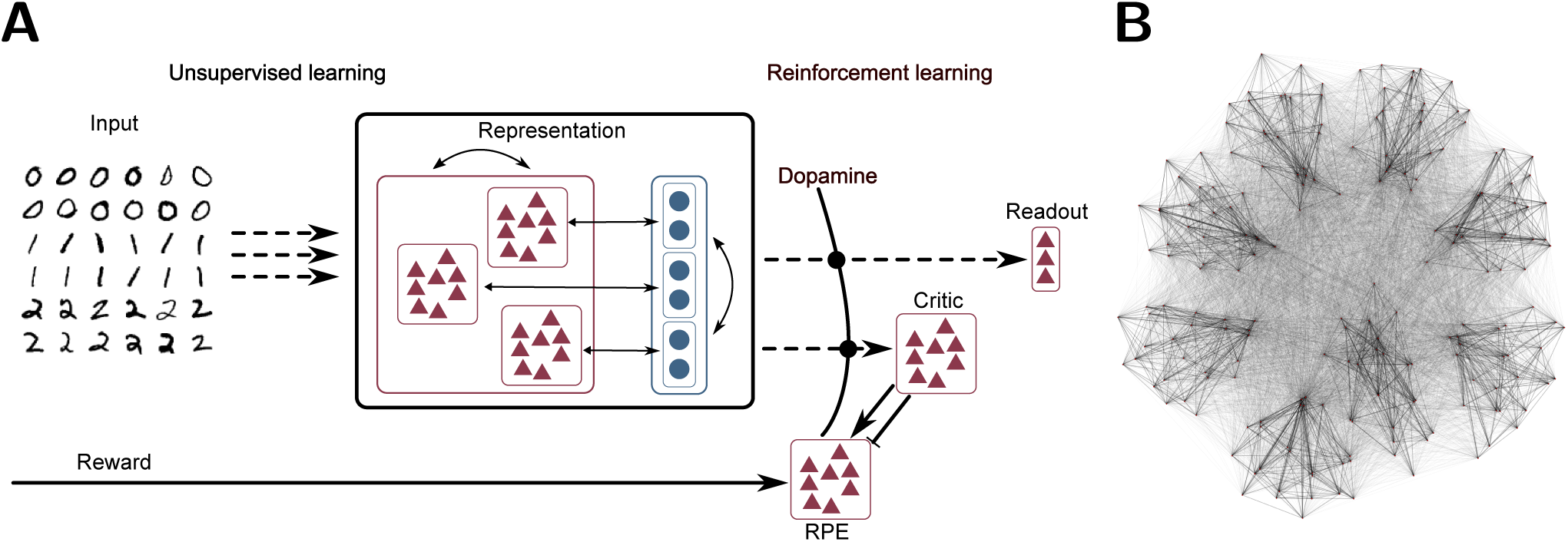
**A** Network schematic. Dashed lines represent plastic synapses. Synapses between the input layer and the representation layer are trained in an unsupervised fashion, synapses to the critic and readout are trained with reinforcement learning. **B** Visualization of network connectivity of a subset of the clustered balanced random network.

#### Input layer

The input layer is a population of rate modulated Poissonian spiking neurons with a maximum firing-rate of *F*_max_. It converts the analog input data into spike trains for the representation layer. For example, a gray-scale image can be presented to the network by stimulating each input neuron with the gray-value *g*_*i*_ of a specific pixel in the image, resulting in Poissonian spike trains with a rate 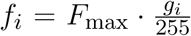. See Section 2.3 for details on the input conversion of each task.

#### Representation layer

The input layer projects to the representation layer, consisting of 5000 integrate-and-fire neurons of which 4000 are excitatory and 1000 are inhibitory. This is implemented as a balanced random clustered network, as described in (Rost et al., 2018; Rostami et al., 2020): the connection probability is uniform, but synaptic weights within a cluster (containing both excitatory and inhibitory neurons) are scaled up while the weights between clusters are scaled down, such that the total synaptic strength remains constant (see Figure 1B for a visualization of the network). Thus, the neurons in the representation layer receive input from the input layer and from recurrent connections:

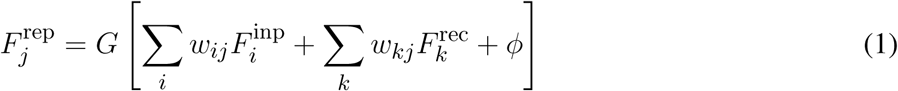

where *G* is the non-linear transfer function of the integrate-and-fire neuron and *ϕ* is a static background input (bias).

#### Output layer

The output layer is realized by a soft winner-takes-all circuit of integrate-and-fire neurons, i.e. the number of output neurons corresponds to the number of labels in the dataset (for classification tasks) or the number of actions (for reinforcement learning tasks with discrete actions) and, at any given point in time, only one output neuron is active, i.e. the neuron with the highest weighted input:

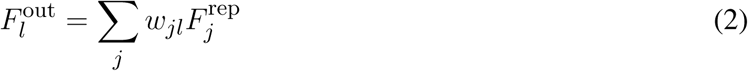

The predicted label *ŷ* (or chosen action) is then defined as the most active output neuron 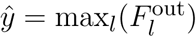.

#### Actor-critic circuit

In discrete time/state temporal difference learning, the reward prediction error is calculated from the expected value *V* of two successive states and the reward received for the last action (if any):

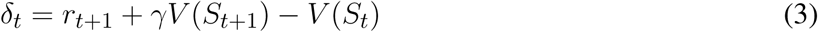

where *γ* is the discounting factor. For values of *γ* close to zero, the agent is short-sighted and prefers immediate reward to rewards in the future. Values close to one correspond to a strong weighting of future rewards. This RPE is then used to update the expected value of the previous state and the policy of the agent, such that actions leading to better states than expected become more likely, and vice versa. In the context of neural activity, Frémaux et al. (2013) showed that a continuous signal, associated with the concentration of dopamine, can play the role of a reward prediction error:

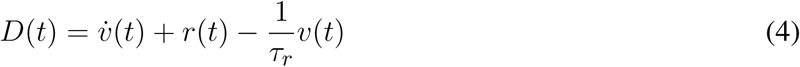

where *v* is the rate of a critic neuron (or population of neurons), *r* is the reward received directly from the environment and *τ*_*r*_ is the discounting time constant.

We implement the critic as a population of 20 Poissonian spiking neurons. The rate of this population is taken as an approximation of the value of the current active cluster and projects to a population of 1000 Poissonian spiking neurons representing the reward prediction error (RPE), which in turn produce the dopaminergic concentration *D*(*t*) as described above. The instantaneous change of 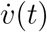 is implemented (as in Jordan et al. 2017) as a double connection from the critic to the RPE where one connection is excitatory with a small delay of 1 ms and the second is inhibitory with a larger delay of 20 ms (Potjans et al., 2009; Jitsev et al., 2012). Note that no claim is made for the biological plausibility of this circuit; it is simply a minimal circuit model that generates an adequate reward prediction error to enable the investigation of the role of clustered structure in generating useful representations for reinforcement learning tasks. The RPE signal enters the plasticity of the synapses between the representation layer and the output layer (i.e. the actor) as a third factor, as described in the next section.

### 2.2 Plasticity

The projections from the input to the representation layer are subject to an unsupervised, local plasticity rule, as proposed by Tetzlaff et al. (2013):

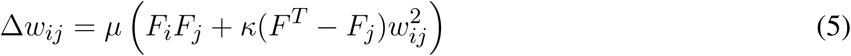

The learning rule comprises a Hebbian component, dependent on pre- and post-synaptic activity (*F*_*i*_ and *F*_*j*_, respectively) and a non-Hebbian, homeostatic term with a quadratic weight dependence (see Tetzlaff et al. (2013) and references therein). While the Hebbian term establishes an appropriate, input-specific mapping, the homeostatic term guarantees convergence and stability in the resulting weights, while dynamically retaining their integrity (relative weight distribution). The parameter *µ* sets the global learning rate, whereas *κ* controls the ratio of synaptic scaling relative to Hebbian modifications. Neurons’ activations *F* are calculated by low-pass filtering the spike trains with an exponential kernel with a fixed time constant *τ* = 100 ms and *F* ^*T*^ constitutes the *homeostatic set point*, i.e. the target firing rate.

The synapses between the input and representation layers evolve according to an unsupervised local learning rule, as described in Section 3.1 and Equation (5). This rule is extended for the synapses from the representation to the output layer to incorporate an explicit reinforcement signal (making it a three-factor learning rule). For these synapses, the Hebbian term in Equation (5) is complemented with a multiplicative modulatory term. Learning in the output projections thus takes the form:

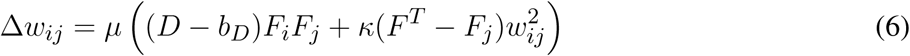

The reinforcement signal is modelled as the dopaminergic concentration *D* relative to its baseline value *b*_*D*_, which in turn is computed as a moving average over the last 10 s.

This segregation between purely unsupervised learning in the input projections versus reinforcement learning in the output projections is illustrated in Figure 1A.

### 2.3 Tasks

#### 2.3.1 MNIST task

MNIST^1^ is a dataset of handwritten digits from zero to nine. Each digit is presented as 28×28 pixel grayscale images and the whole dataset contains 60000 training images and 10000 test images. As the runtime of our network is too slow to present the complete dataset, we use only the first three digits. We further reduce the 18000 training and 3000 test images to 1000 randomly picked images for the training phase and 150 for the test phase.

For each trial during the training phase, one digit was picked randomly and presented to the network. The images are translated into spiking activity using one neuron per pixel as described in Section 2.1. The error signal *D* is generated using a neuron which doubles its base firing rate for 100 ms if the label *ŷ* produced by the network is the correct label *y*, and becomes silent for 100 ms if it is incorrect.

#### 2.3.2 Mountain Car task

Mountain Car^2^ is a reinforcement learning environment of the OpenAI Gym. In this task, a car is randomly placed between two hills. The goal is to reach the top of the right hand hill. This task is difficult because the engine of the car is not strong enough to just drive up the hill in one go; instead the agent must build up momentum by swinging back and forth until the car finally reaches the top (see Figure 2 **C**).

**Figure 2.**
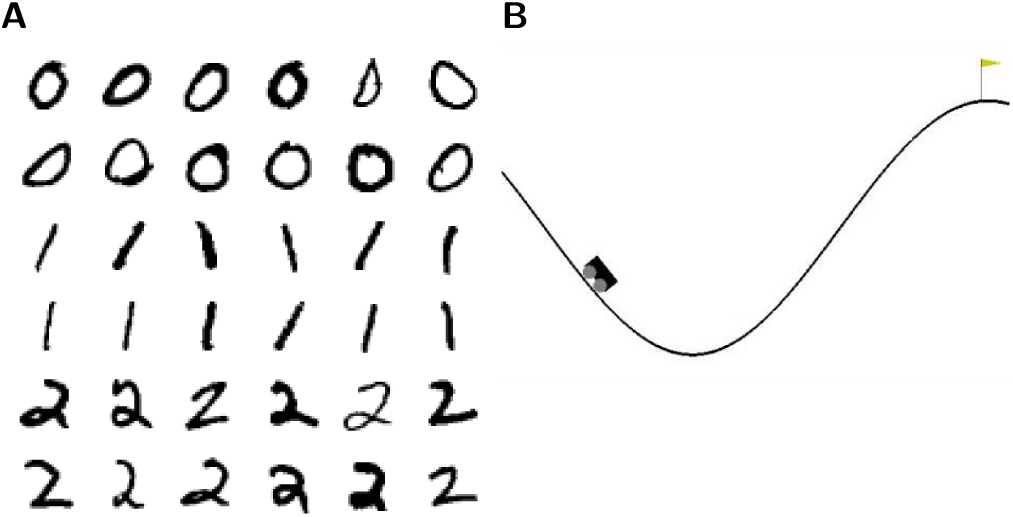
Visualization of the tasks. **A** 36 example images of the 3-digit subset of the MNIST database. **C** Visualization of the Mountain Car environment of the OpenAI Gym.

The available information for the agent consists only of the position and the velocity of the car and, in every timestep, the agent receives a constant punishment of −1. The episode terminates, and the environment resets, if the car reaches the goal position or a limit of 200 timesteps is reached.

We adapted the environment in two minor ways. First, we removed the time limit of 200 timesteps in order to enable our agent to explore the environment. Second, we added a positive reward of 1 when the car reached the goal position to give the reinforcement learning algorithm a higher contrast to the constant punishment during the trials.

The OpenAI Gym defines the task as solved if less than 110 timesteps are needed to reach the goal for 100 consecutively trials. This is not easy to achieve as the car starts in a random position between the hills and in some cases it is necessary to first swing to the right, then to the left and finally to the right again. From the worst starting position the timing must be perfect to solve the task within 110 timesteps.

The input is presented to the network by representing each input channel (position, velocity) with *N* = 200 spiking neurons emitting spike trains with Poissonian statistics. Each channel is normalized between −1 and 1 and each neuron *i* within one channel has a Gaussian shaped receptive field of mean 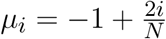 and sigma *σ*_*i*_ = 0.05. The activation of the neurons are translated to Poissonian spike trains with a firing-rate between 0 and 35 spks*/*s

The readout consists of three neurons representing the three different actions ‘left’, ‘right’ and ‘nothing’. As in the other tasks, the readout neurons are in competition, ensuring that they are not active at the same time.

### 2.4 Simulation tools

All neural network simulations were performed using the Neural Simulation Tool 2.16 (NEST) (Gewaltig & Diesmann, 2007; Linssen et al., 2018). The interface between NEST and the OpenAI Gym (Brockman et al., 2016) was implemented using the ROS-MUSIC Toolchain (Weidel et al., 2016; Jordan et al., 2017). Simulation scripts and model definitions are publically available ^3^.

## 3 RESULTS

### 3.1 Learning input representations and resolving non-linearities

The computational benefits conferred by the ability to learn suitable input representations are clearly demonstrated by the simple example shown in Figure 3, employing the logical operation ‘exclusive- or’ (XOR), which is the most fundamental non-linear task. For this example, we define an *input layer* comprising two neurons (A and B) which represent ‘1’ and ‘0’ by being either highly active or nearly silent, respectively. The activity of these input neurons is randomly projected to a higher-dimensional space spanned by twelve integrate-and-fire (IAF) neurons forming a weak winner-take-all network, the *representation layer*. This simple setup is illustrated in Figure 3A.

**Figure 3.**
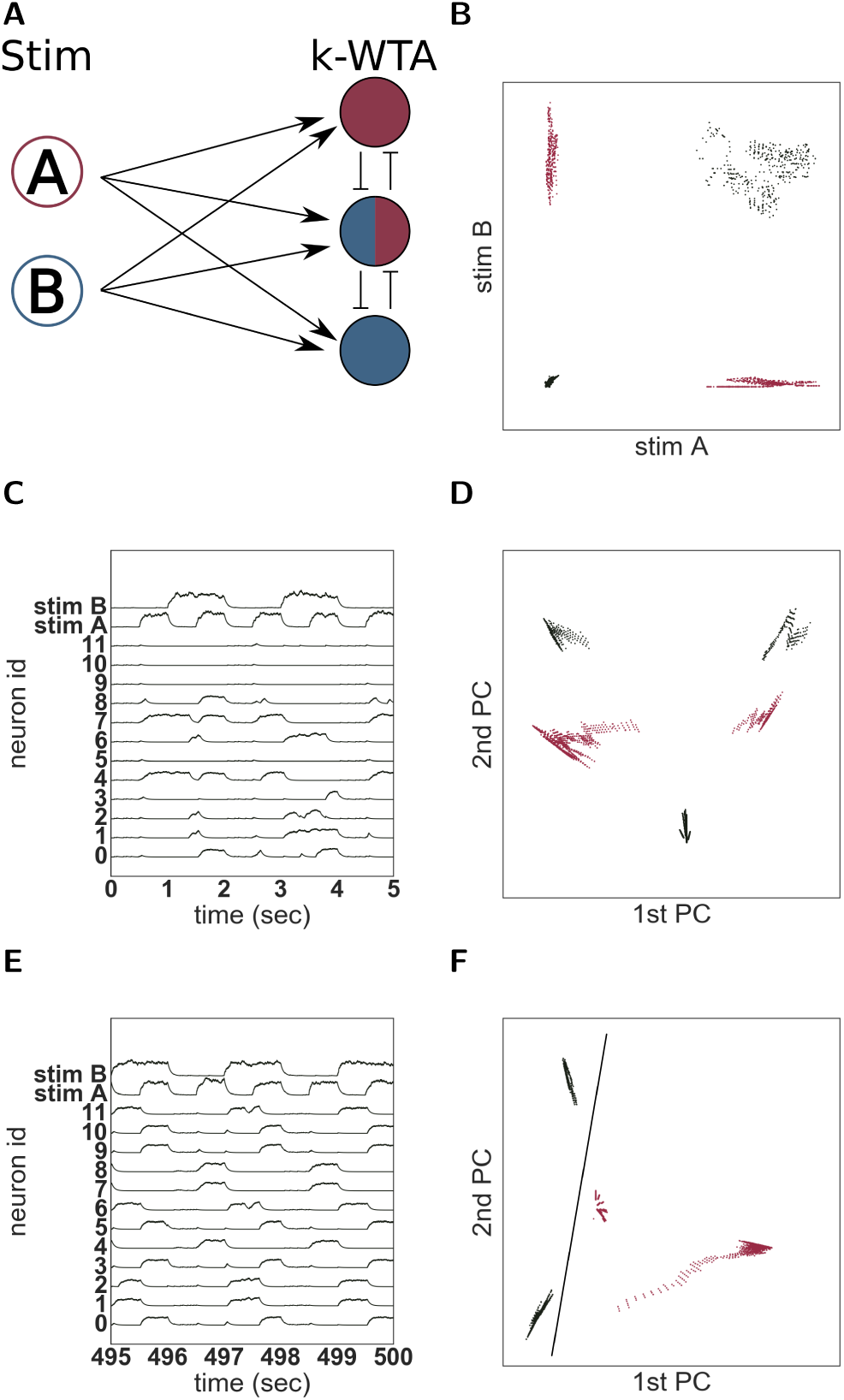
Solving the XOR task with unsupervised learning. **A** Network schematic of a two-layered network to solve the XOR task comprising two stimulation neurons (A and B) and twelve representation neurons (of which three are illustrated) implementing a k-winners-take-all architecture. **B** Input activity with black dots representing stimulus patterns ‘00’ and ‘11’ and red dots representing ‘01’ and ‘10’. **C** Neural activity before training. **D** The first two principal components of the first five seconds of activity. **E** Neural activity after training. **F** First two principal components of the last five seconds of activity.

The activity of the input neurons cannot be linearly separated to represent XOR(A, B) as shown in Figure 3B. Whereas the XOR task can be trivially solved by trained artificial neural networks (Gelenbe, 1989) and even by untrained recurrent networks of varying degrees of complexity (using the reservoir computing approach, e.g. Haeusler & Maass 2007; Verstraeten et al. 2010; Zajzon et al. 2019), the combination of a small network size and random input projections chosen in this illustrative example results in an inadequate transformation that still doesn’t allow the nonlinear task to be resolved. This can be seen in the activity of the neurons in the representation layer (Figure 3C), some of which are silent for all stimuli while others are active for multiple stimul. In other words, due to the random input projections, the neurons show no input specificity. Consequently, the two classes are not linearly separable on this neuronal representational space, as can be illustrated on the basis of the first two principal components of the population activity (Figure 3D).

However, we show here that even this small and simple network can resolve the non-linearities in the input, if it can adapt the projection of the input signal onto the neuronal representational space, in order to develop clear stimulus representations. In this particular case, an ideal mapping between stimuli and active neurons would be bijective, i.e. one stimulus should always activate the same set of neurons and one particular neuron should only be activated by one stimulus (exclusive tuning).

To support the reliable formation of such stimulus-tuned neurons and maximize separability, the projections from the input to the representation layer are subject to an unsupervised, local plasticity rule with Hebbian and homeostatic components (see Equation (5) and Tetzlaff et al. 2013). The Hebbian part of the plasticity rule strengthens the weights between stimuli and active neurons and increases the probability that the same input evokes the activation of the same neurons. The scaling part introduces competition between the neurons and ensures that an active neuron is not becoming active for all stimuli. The plasticity rule has some similarities to Oja’s rule (see Oja 1982) which is known to be able to learn a PCA transformation in a similar network setup of rate neurons.

The consequences of this unsupervised rule can be clearly seen by comparing the activities of the neurons in the representation layer before and after learning (Figure 3C, E). After a few hundred seconds of simulation, the neurons show clearly separate receptive fields and are active for only one of the four different stimulus configurations. The projection of the low-pass filtered activity of the twelve neurons onto a two dimensional PCA space (Figure 3F) reveals that their activity is easily linearly separable; thus, the network would now be capable of solving the XOR task. This is unsurprising given the known effects of this type of plasticity rules (Tetzlaff et al., 2013), but it demonstrates that learning introduces important specificities into the population dynamics that allow it to be used as a substrate for further computation.

### 3.2 Feature extraction in a classification task

In the previous section, we demonstrated that the unsupervised learning rule we implement is capable of learning input representations that enable a simple system to resolve non-linearities in the input signal. To further examine its capabilities, we now explore a more challenging task and a more complex system. The task we explore in this section is handwritten digit recognition (the MNIST dataset), most commonly used for image classification, for which the algorithm has to detect handwritten digits in grayscale images of size 28 × 28. Due to the high computational load for the numerical simulations, we reduce the full dataset to a three digit subset. Moreover, we limit the training set to 1000 images and the test set to 150 images, which is just a small fraction of the available data. Details of the experimental set-up are given in Section 2.3.1.

The network structure is illustrated schematically in Figure 1A and described in detail in Section 2.1. As for the previous task, the network we implement consists of an input and representation layer, but is now supplemented with an *output layer*, which is realized by a soft winner-takes-all circuit of three integrate-and-fire neurons, one for each label in the dataset; the most active neuron is interpreted as the network’s decision on which label corresponds to its current input.

As the input dimensionality is much larger in the MNIST task than in the logical XOR task, a larger representation layer is needed, which we implement as a clustered balanced network as described by Rost et al. (2018). Unlike the clustered network architecture investigated by Litwin-Kumar & Doiron (2012), we cluster both excitatory and inhibitory connectivity. Rost et al. (2018) showed that networks comprising purely excitatory clusters tend to fire at saturation in the active clusters, with very low activity elsewhere, resulting in only infrequent switches between active clusters. By introducing inhibitory clusters, the firing rate of the active cluster does not saturate, which allows the clusters to switch more easily.

For the purposes of comparison, we consider both a network parameterized to have eight clusters (visualized in Figure 1B) such that the competition between the clusters is very strong and a switch between the winning clusters is rare, and an equivalent balanced network with random connectivity, i.e. no clusters. The total amount of excitation and inhibition in both networks are identical, which is reflected in their identical average firing-rate of ∼27 spikes per second. Dynamically, the difference can be measured using the coefficient of variation (CV) of the inter-spike-intervals. The average CV of the unclustered network is 0.92, which is close to the expected value of a Poisson process. In contrast, the neuronal activity in the clustered network varies much more (CV 4.61) due to the fluctuating activity of the clusters.

The synapses between the input and representation layers evolve according to an unsupervised local learning rule as for the XOR task, and the synapses between the representation and output layers adapt according to an unsupervised rule with an additional neuromodulatory multiplicative term, see Equation (5) and Equation (6). Thus, as illustrated in Figure 1A, the input projections are subject to purely unsupervised learning, whereas the output projections, due to the neuromodulatory third factor, are subject to reinforcement learning.

Note, firstly, that the reinforcement learning signal presented to the output projections denotes only success or failure (see Section 2.3.1) and is therefore less informative than the supervisory learning signal usually used for this task (which would denote the correct choice in case of failure), and secondly, that there is no error back-propagation from the output layer to the representation layer to help to solve the credit assignment problem. Thus, the classification performance depends on a stable and consistent representation of the input.

#### 3.2.1 Self-organisation of feature extracting representations

Figure 4**A** shows the evolution of spiking activity in the unclustered network for one trial of the MNIST task. The neurons are ordered by their maximal response to the three different input classes; the excitatory neurons most responsive to stimuli of class *‘zero’* are at the bottom of the plot, then those most responsive to digit *‘one’*, followed by digit *‘two’*. This ordering is repeated at the top of the plot for the inhibitory neurons. In the first five seconds (left panel), no clear distinction can be made between the activity of the network on the basis of the input digit, although some overall changes in firing rate can sometimes be observed between two input stimuli. The white horizontal bar in the plot is due to neurons that do not fire at all during the recording period. However, by the last five seconds (right panel), there are easily discernible differences in the spiking activity corresponding to the individual input digits. Consistent with the results presented in Auth et al. (2018), employing a similar learning rule, the network has self-organized into effective clusters - effective in the sense that only the input weights have adjusted, not the recurrent weights - that represent the digits *‘zero’, ‘one’* and *‘two’*. The effect of this self-organization on the network dynamics is most evident in the coefficient of variation of the spike trains, which increase from 0.9 at the beginning of the trial to 2.7 at the end of the trial. The average firing rate is hardly changed, dropping from 27 to 26 spikes per second.

**Figure 4.**
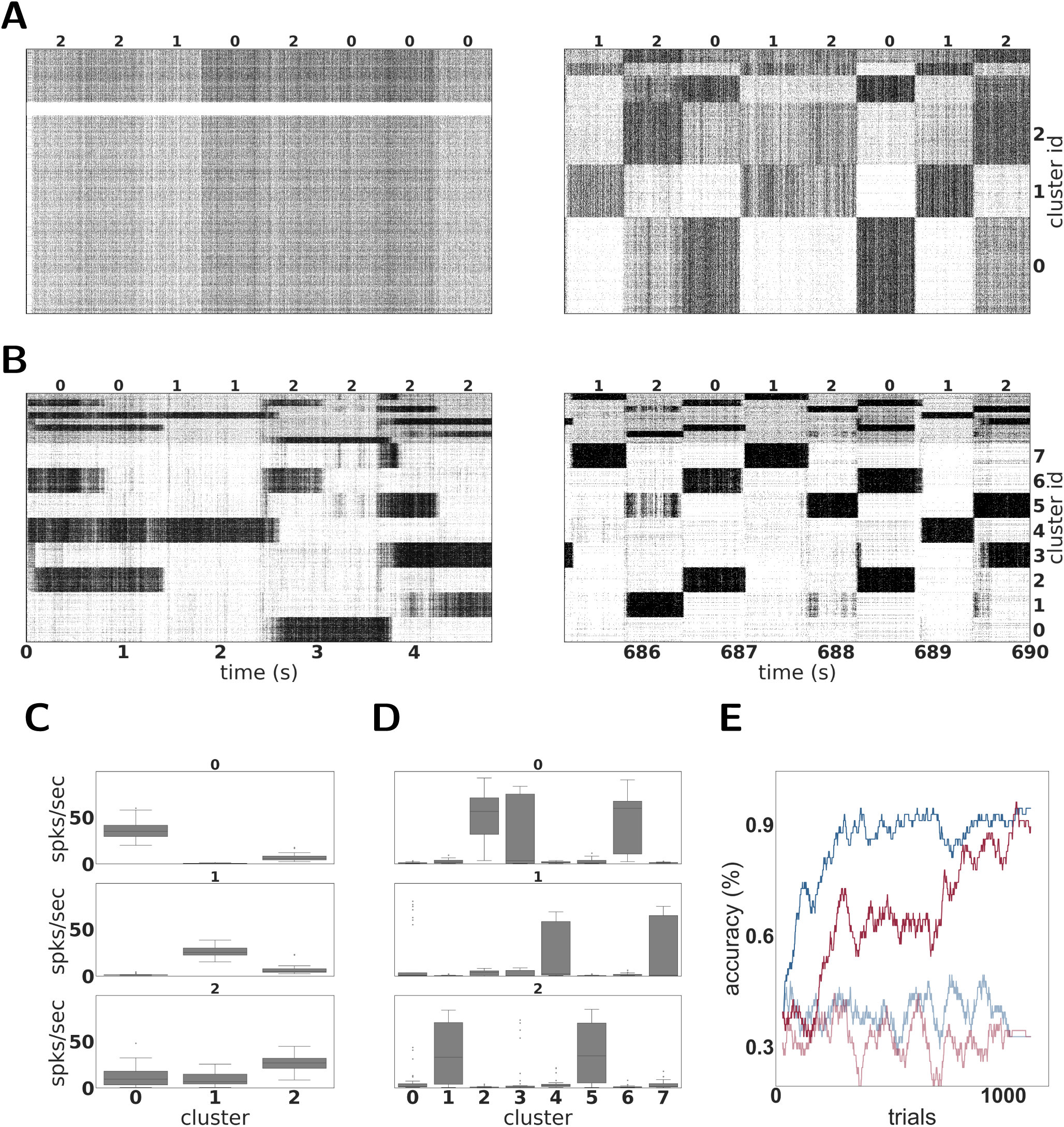
Dynamics and performance of the model during the MNIST task. **A** Raster plot of an unclustered balanced random network model during the first and the last five seconds of the simulation while solving the MNIST task. The identity of the stimulus digit is shown above the plots. The first 4000 neurons are excitatory, and the last 1000 are inhibitory. Some neurons are silent in the beginning of the simulation. **B** As in **A**, but for a network with eight clusters. **C** Activation of the three emergent clusters in the initially unclustered network during the presentation of the last 100 stimuli (indicated above the panels). **D** As in **C** but for the network with eight clusters. **E** Classification performance of the unclustered (red) and clustered (blue) network (one instance each). Curves in pale colors indicate the performance of the corresponding networks without unsupervised plasticity on the input projections. Performance is calculated using a sliding average over a window of 30 trials.

In contrast to the unclustered network, the activity in the clustered network is already strongly differentiated in the first five seconds (see Figure 4**B**, left panel). The clusters switch on and off at the transition points between stimuli. At this point, it is hard to discern stimulus-specific patterns - for example, cluster 0 is sometimes on for a presentation of the digit *‘two’*, and sometimes off. By the end of the trial (right panel), the activity has coalesced into clear stereotypical patterns: as an example, stimuli from the class *‘one’* always evoke activity in clusters 4 or 7. The change in the network response to stimuli is less visible on the level of spike train statistics than that of the unclustered network; the CV increases from 4.6 to 5.2 and the average firing rate decreases from 27 to 25 spikes per second.

The detailed response profiles for each stimulus class can be seen in Figure 4**C,D**. Notable here is that, while each stimulus can be represented by more than one cluster, there is no overlap between the profiles - each cluster responds to only one stimulus class.

The performances of the two network configurations are shown in Figure 4**E**. In the absence of unsupervised plasticity at the synapses between the input and the representation layer, both the unclustered and the clustered network perform at chance level. In the presence of plasticity, both networks reach a good performance, clearly demonstrating the importance of learning input projections for this task. Of the two networks, the clustered network performs consistently better - it learns faster, reaching an accuracy level of 80% after 181 trials and 90% after 268 trials, compared to 786 and 1014 for the unclustered network. The performance of the clustered network is also better after saturation than that of the unclustered network. Over the last 100 trials, the clustered network has an average of 96%, compared with 92% for the unclustered network. For perspective, the current best performance on the entire MNIST dataset was achieved by a committee of convolutional neural networks (∼99.7%^4^). Thus, we consider the performance of the spiking networks satisfactory, considering the small fraction of data used for training and the use of a reinforcement learning signal rather than the more informative supervisory feedback.

To acquire some insight into why the clustered network performs better, we consider the features of the input data and their representation in the network. The three digits selected from MNIST have a large overlap and are not linearly separable, similar to the XOR task examined in Section 3.1. However, in contrast to XOR, the classes in MNIST have a large intra-class variance: there are sub-classes of different styles of writing the digits. To give some examples, there is a sub-class of the digit *‘zero’* written close to circular while other sub-classes are more tilted or oval; the digit *‘one’* is often written as vertical line, but sometimes also tilted; the digit *‘two’* may be written with or without a loop. A sample of these varying styles can be seen in Figure 2**B**. This intra-class variability is part of what makes the MNIST task challenging. In order to solve the task, a classifier needs to learn a category that is broad enough to detect all the instances of a particular digit, whilst simultaneously being narrow enough to exclude all the instances of other digits.

The mean synaptic weights between the input neurons and the neurons in each (effective) cluster of the representation layer corresponds to the receptive field of each cluster, and thus the specialization learned in an unsupervised fashion by that cluster. These can be visually compared in Figure 5. The receptive fields of the effective clusters that self-organize in the unclustered network are generic for each digit (Figure 5**A**). This results in a blurred appearance, because they contain traces of all the sub-classes of a particular digit. In contrast, the receptive fields of the clustered network have a less blurred appearance and reveal that each cluster has typically specialized itself for a specific sub-class. Thus, there are different clusters representing a round or oval *‘zero’*, a straight or tilted *‘one’*, and a looped or unlooped *‘two’*. This specialization explains the pattern of activity observed in Figure 4**B**, right panel: different sub-classes of a digit evoke activity in different clusters. In addition, this explains the wide variance of the response profiles shown in Figure 4**D** and the performance shown in Figure 4**E**: the more specific receptive fields learned by the clustered network allow a stronger differentiation of a cluster’s response to its preferred digit over non-preferred digits than the generic receptive fields of the unclustered network, leading to a higher accuracy.

**Figure 5.**
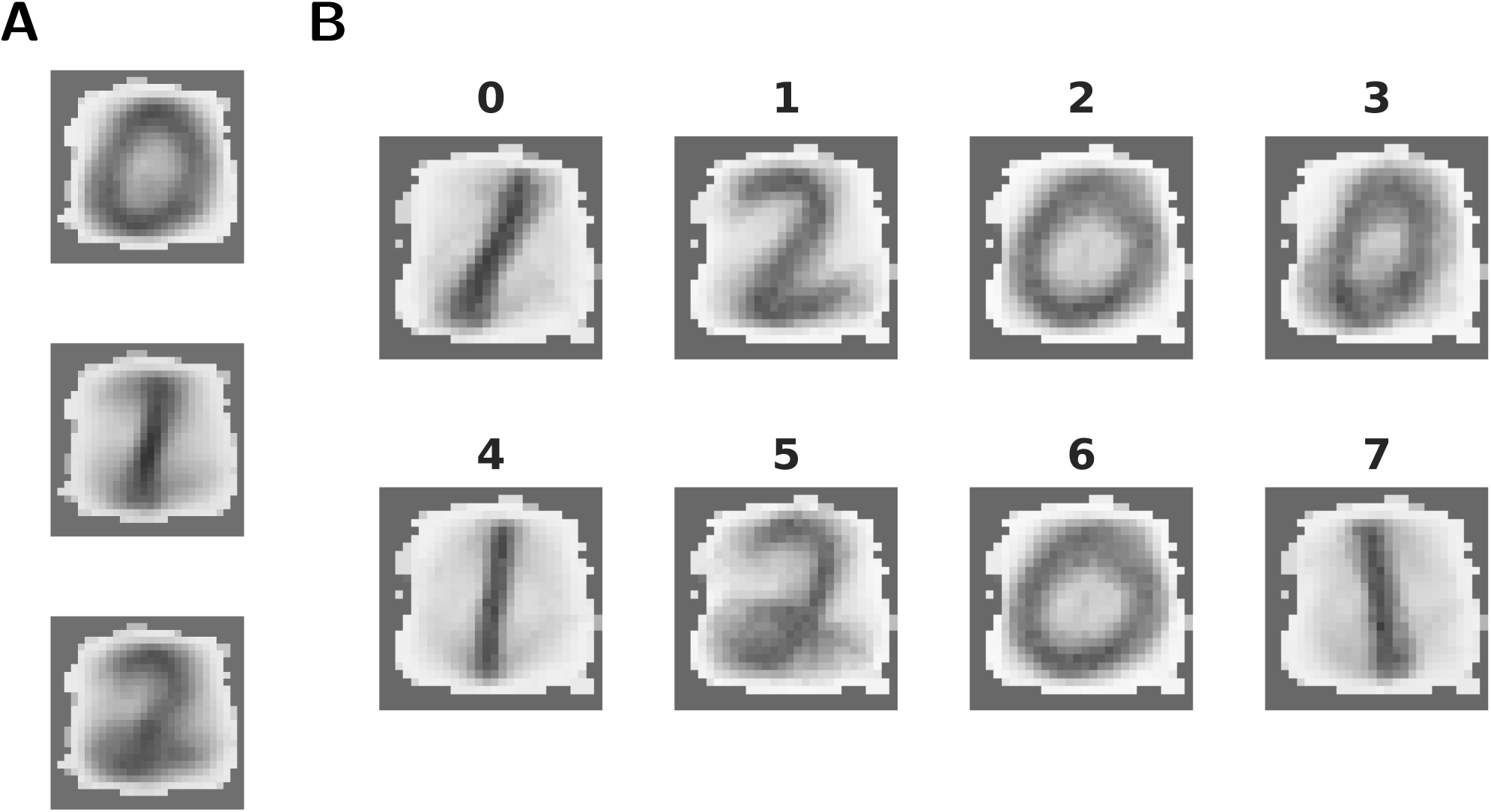
Features extracted by unsupervised learning. **A** Mean synaptic weights between the 28 × 28 input neurons and the three effective clusters of the initially unclustered network, see Figure 4A for the corresponding activity. **B** As in **A**, but for the eight clusters of the clustered network (cf. Figure 4B).

We conclude that the pre-existence of clusters supports the self-organization of representation by permitting feature specialization. The specialized classifiers learned by the cluster thus allow a faster and more robust learning performance. Note that this specialization is an emergent property of the combination of the clusters and the synaptic plasticity between the input and the representation layers, and not of the task. The evolution of these synaptic weights is independent both of the label of the data and the prediction error, and depends only on the structure of the input data.

### 3.3 Continuous time and space and delayed and sparse reward

So far we have shown that our model can resolve non-linearities in the input and extract and represent complex features in images using the XOR and the MNIST classification tasks. In both cases, to train the readout weights with reinforcement learning, the reward or punishment is applied directly after the end of the stimulation period. This is not a very representative scenario for real-world tasks, where it is necessary to make a sequence of decisions which typically involve a substantial delay period until the reward is received (Tervo et al., 2016; Schultz, 2016). Moreover, unlike XOR and MNIST, which constitute toy examples, real-world stimuli unfold continuously in both time and space. Foraging is a prime example of a real world task which has the properties of continuous state-space and delayed reward.

We therefore investigate the performance of our model in a more challenging task, continuous in time and space and providing only sparse and delayed reward at the end of a successful trial: the ‘Mountain Car’ reinforcement learning environment provided by the OpenAI Gym. In this task, a car is placed in the valley between two hills. The task is solved when the agent learns to drive up one of the hills by swinging back and forth to gain momentum (see Section 2.3.2). The only information available to the learning agent is the continuous position and velocity of the car.

As the reward is only presented at the end of a trial, we include an actor-critic architecture in our network structure. This is illustrated schematically in Figure 1**A** and described in detail in Section 2.1.

The actor-critic approach is commonly used in reinforcement learning tasks where an immediate reinforcement signal is not available (Sutton & Barto, 2018). In this framework, the actor selects the actions, whilst the critic calculates the expected value (i.e. the discounted total expected future rewards) of the environment’s states. From the difference between the value of the state before the action, and the sum of the reward received for an action and the discounted value of the new state, a reward prediction error (RPE) can be derived which expresses whether the action selected produced better or worse results than expected. This signal can then be used to update both the expectations of the critic and the policy of the actor.

Generating a RPE allows a learning agent to project a sparse reward back to earlier (non-rewarded) states in the episodes. However, doing so requires a stable representation of environmental states. Whereas Frémaux et al. (2013) was able to show that the continuous time neuronal formulation of the RPE given above can solve a variety of tasks including maze navigation and dynamic control, and Jordan et al. (2017) later also demonstrated that this rule is sufficient to solve the Mountain Car problem investigated here, both of these studies depended on a hard-coded ‘place cell’-type representation of the environment, as indeed is typical for models of non-trivial reinforcement learning in neuroscience (e.g. Potjans et al. 2011; Frémaux et al. 2013; Friedrich et al. 2014; Jordan et al. 2017).

In our model, however, an appropriate representation of the environment is automatically learned by the clusters of the BRN. After exploring the environment for long enough, the clusters in the BRN become active for those parts of the state space which are frequently visited and therefore form a task-oriented, rather than comprehensive, discretization of the state space. The mapping of clusters to portions of the two dimensional task space can be clearly seen in Figure 6**A**. Due to this mapping, the clusters are activated in a stereotypical sequence, as is visualized as a directed graph for a specific instantiation in Figure 6**B** and is also readily apparent in the spiking activity displayed in Figure 6**C**. For this instantiation, clusters 6 or 2 represent the starting positions and are therefore activated first in the sequence. Cluster 4 represents the goal position and is activated last in the sequence.

**Figure 6.**
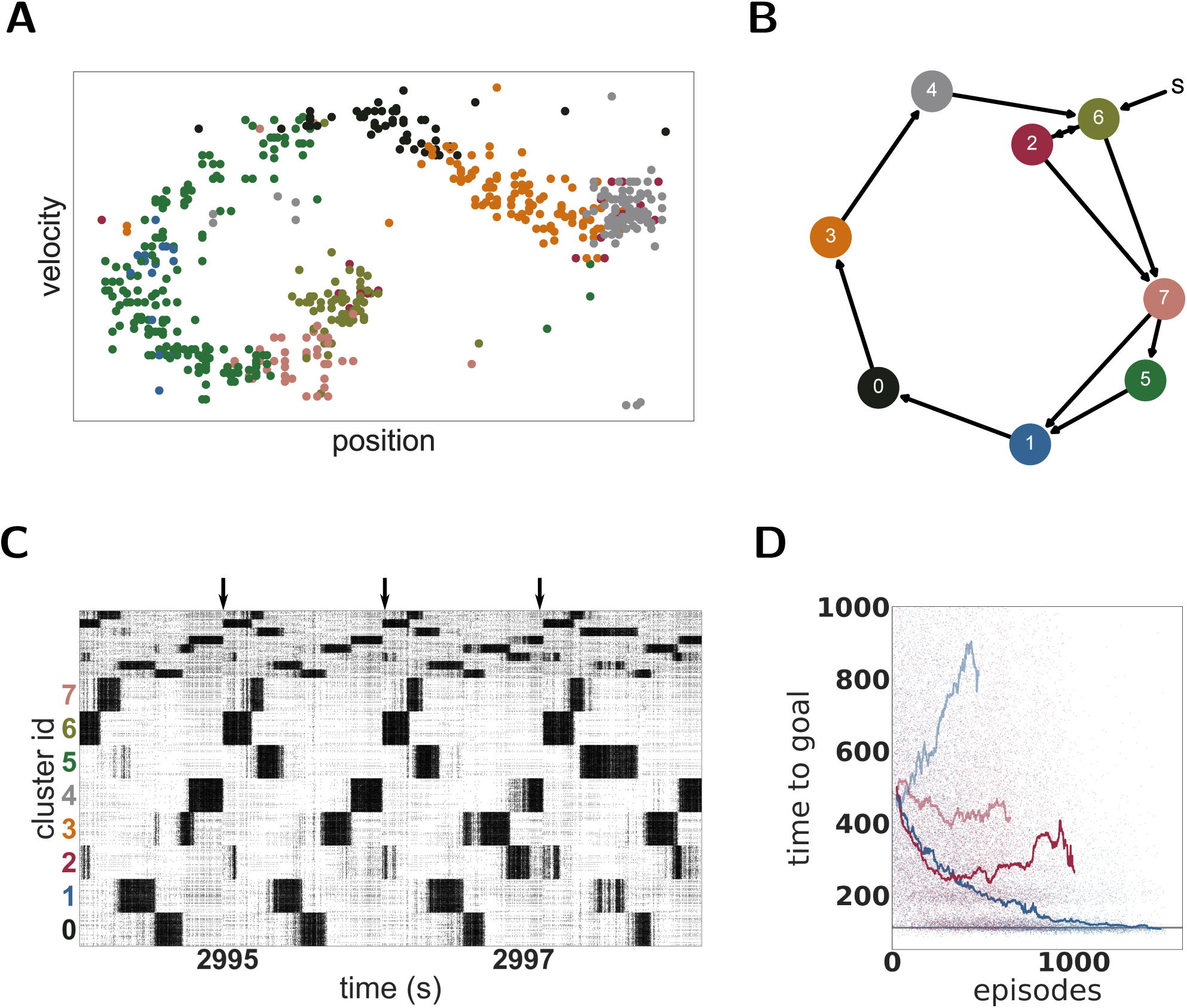
Dynamics and performance of the model during the Mountain Car task. **A** Each dot represents the reconstructed position and velocity of the car in the input space during one trial (corresponding to the last trial of panel D). The color of the dots correspond to the most active cluster during this velocity-position pair. **B** Most common sequence of active clusters after training. The sequence of each trial usually starts with cluster 6 or 2 and ends on cluster 4. **C** Raster plot of the last seconds of the simulation of the Mountain Car task. Arrows indicate the approximate start of a new trial. **D** Each dot represents the time to reach the goal in one episode. Curves give the running median over ten runs of the model for the unclustered (red) and clustered (blue) networks. Dark and pale curves indicate plastic and static input projections, respectively.

The emergent representation of the environment in the BRN provides a basis for the actor-critic architecture to evaluate states and learn an effective policy. OpenAI Gym describes the task as solved if the agent needs less than 110 timesteps to find the goal in 100 consecutive episodes. Our model approaches this threshold within 1000 episodes (see Figure 6**D**) but for two reasons we can not claim to have solved the task. First, we adapted the environment in two minor ways as described in Section 2.3.2 in order to give our agent more time to explore the environment. Second, although our model frequently finds the goal in less than 110 timesteps, it does not do this consistently. The model is constantly exploring, learning and changing its policy which results in a substantially sub-optimal result in a few trials. Over the last 100 trials, it found the goal in under 110 timesteps on 59 trials, with a median of 108 and an average of 164 timesteps, respectively. As a side remark, humans cannot solve the task easily either. Our personal experience revealed that reaching the goal 100 times in a row under 110 timesteps was challenging and tiring.

As with the MNIST task, it is the combination of a clustered network and plastic input synapses that produces the best performance. As shown in Figure 6**D**, an unclustered network with plastic synapses shows initial improvement on the task but saturates early at just over 200 timesteps, and then becomes gradually worse (median 329, average 416 timesteps over the last 100 episodes). An unclustered network with static input synapses does not show appreciable change in performance even after many iterations (median 437, average 541). Unlike the MNIST task, for the Mountain Car task the worst performance is demonstrated by the clustered network with static synapses (median 844, average 1086). In this configuration, cluster switching becomes rare, and so the network takes longer and longer to reach the goal.

Also in common with the MNIST task, the change in coefficient of variation is the best indicator of performance. For the networks with plastic input synapses, the CV of the clustered network increases during the simulation from 0.43 to 3.99 (whilst the firing rate increases mildly from 15.5 to 18.8 spikes per second); the CV of the unclustered network increases from 0.93 to 2.08 (whilst the firing rate increases substantially from 8.8 to 22.8 spikes per second). For the networks with static input synapses, the CV of the unclustered network stays constant at 0.78 whereas the CV of the clustered network decreases from 0.69 to 0.55.

## 4 DISCUSSION

In this work, we present a spiking neural network model for unsupervised feature extraction and reinforcement learning using clustered spiking networks and Hebbian plasticity with synaptic scaling. We demonstrate that this combination is able to extract complex features and resolve non-linearities in the input elegantly. The XOR task (section 2.1) demonstrated that the presence of unsupervised plasticity on the input projections can boost the performance of a reservoir computing system, even in a very small and simplified network. Whereas this simple network with static input projections was unable to transform the input into a linearly separable projection required to resolve the task, simply adding plastic, unsupervised input projections allow it to adequately do so.

Using the full version of the model, with a balanced random network as the representation layer, we showed that the combination of unsupervised learning on the input projections with clustered connectivity boosts the performance of a reinforcement learning approach. On the reduced MNIST task (Section 3.2), the clustered network with unsupervised plasticity learns faster and achieves a better performance than an unclustered network. Networks without input plasticity performed at chance level. Examining the structure of the input projections revealed that the clusters had specialized for particular sub-categories of the input space (e.g. ‘*two*’s with and without a loop, see Figure 5).

A spiking implementation of an actor-critic reinforcement learning architecture enables the model to solve tasks with sparse and delayed feedback, such as the time- and space-continuous Mountain Car problem. On this task, the clusters became specialized for particular regions of the two dimensional input space, for which the appropriate action could easily be learned. Thus, the model cycled through a stereotypical sequence of cluster activations in order to solve the task, and achieved a much better performance than the corresponding unclustered network (see Figure 6).

Through the effect of a local and entirely unsupervised learning rule, compatible with biological findings, the model is thus able to learn suitable, task-relevant input projections that lead to stable and linearly separable representational states. The pre-existence of clusters supports the self-organization of the representations by allowing neuronal specialization to develop, i.e. if the input is adequately projected onto clusters of more strongly connected neurons, competition between the clusters allows them to become tuned to the relevant input features driving them, such that the system learns task-relevant mappings faster and in a more robust manner. Importantly, this specialization develops independently of the task feedback that drives learning (labels, errors, etc.), relying only on the statistical structure of the input data. In the more complex Mountain Car environment, where the partition of the input space is not as clear as in the simple MNIST classification task, the network learns an appropriate partition that reflects the task structure. Cluster activation in this setting reflects the frequency and temporal order with which different regions in the input space are visited, thereby reflecting a task-oriented, rather than comprehensive (grid-like), discretization of the state-space (see Figure 6).

The *representation layer* is left untrained. This has two important ramifications. First, the representation layer does not reflect any previous knowledge or assumptions about an appropriate partitioning of the input space on the part of the modeller, unlike previous approaches with hard-wired place cells or radial basis functions. Second, the network can be re-used to represent different patterns and data sets. As long as the input projections are adapted to the properties of the data, no further modifications are required within our set-up; the exact same network is used for both the MNIST and Mountain Car tasks requiring no modifications of its internal architecture.

The ability to adequately represent data is a critical step for any learning system to be able to detect statistically repeating patterns. Depending on the task and data set considered, it might be important, for example, to extract features that allow a scale- or rotation-invariant representation or to reduce the input dimensionality making it simpler to process. The most commonly known method for dimensionality reduction and feature extraction is principal component analysis (PCA), but many others exist, such as linear discriminant analysis (LDA, Mika et al., 1999), independent component analysis (ICA, Comon, 1994; Hyvärinen & Oja, 2000) or scale-invariant feature transform (SIFT Lowe, 1999). Modern techniques based on convolutional deep networks (CNN, e.g. Mnih et al., 2015) have also proven valuable for feature extraction, particularly in the domain of computer vision, whereby complex, hierarchical feature dependencies are gradually extracted through error back-propagation.

All these methods are powerful tools, but their biological plausibility is limited or non-existent. As such, they are not appropriate models for studying learning in the mammalian brain. A more plausible approach is based on “competitive learning” as described in Hertz et al. (1991), which employs Oja’s rule (modified Hebbian learning with multiplicative normalization). This model was shown to perform PCA, constituting a powerful, biologically-plausible alternative for feature extraction and dimensionality reduction (Oja, 1982), see also Qiu et al. (2012) for a detailed review on neural networks implementing PCA or non-linear extensions of PCA. In a recent study, competitive unsupervised learning applied to the lower layers of a feedforward neural network was shown to successfully lead to good generalization performance on the MNIST dataset (Krotov & Hopfield, 2019). However, despite providing further validation to the claim that biological compatibility need not be sacrificed for adequate computational performance, these studies still rely on implausible mechanisms, such as decoupled processing layers (i.e. purely feedforward connectivity) and simplified processing units (sigmoid or rectified linear units).

Alternative, related instantiations have been proposed in the literature. Employing Hebbian learning with synaptic scaling to the internal, recurrent connections was shown to be mathematically equivalent to non-negative matrix factorization (Carlson et al., 2013) and to allow for the development and maintenance of multiple internal memories, without catastrophic interference effects (Auth et al., 2018). On the other hand, applying similar learning rules exclusively to the input projections (as we have), leads to systems that perform operations similar to ICA (Jonke et al., 2017). These examples employ a conceptually similar learning rule in small network architectures (8, 500 and 900 neurons, respectively). In such small networks, single neurons can become highly tuned and influential enough to reliably implement the competition needed by inhibiting the rest of the network. For larger networks, however, a coordinated activation of relatively large sub-populations is necessary to achieve this internal competition. Our model achieves this competition effect as a result of the clustered connectivity (see section 3.2), and thus highlights the functional relevance of structurally constrained recurrent connections.

The existence of the clustered synaptic connectivity investigated in our study is biologically well motivated (Song et al., 2005; Perin et al., 2011), and can emerge from learning in structured environments (Litwin-Kumar & Doiron, 2014; Zenke et al., 2015). It has been shown that it can account for pervasive phenomena in cortical microcircuits, such as a modulation of the timescales of intrinsic firing fluctuations and their variability (Litwin-Kumar & Doiron, 2012), a drop in effective dimensionality of population responses during active processing as well as the emergence and modulation of stimulus-specific metastable states and structured transitions among them (Mazzucato et al., 2016).

Cortical microcircuits are known to rapidly switch among different active states, characterized by markedly different dynamical properties and critically modulating stimulus processing and the fidelity of stimulus representations (Duarte, 2015; Gutnisky et al., 2017). While the majority of the studies on the matter focus on the relation between ongoing and evoked activity and/or trial-to-trial variability (see, e.g. Churchland et al., 2010, and references therein), much less is known about learning-induced changes in circuit responsiveness and cortical states (Kwon, 2018). A common observation is that the onset of a stimulus reduces the variability of the elicited responses (Churchland et al., 2010) by coalescing population activity onto a low-dimensional manifold (Jazayeri & Afraz, 2017; Remington et al., 2018). One would thus expect that during learning, these representational structures become sharper and this specialization would be reflected in a reduction in response variability.

Our results demonstrate an increase in spike-train variability across the population, but, in this situation, this is clearly a result of modular specialization. Tuned sub-populations of neurons (within a cluster) fire strongly for short periods of time and sparsely when the cluster they belong to is inactive. This skews the ISI distributions, greatly increasing its variance and reducing its mean, resulting in a larger CV. However, trial-to-trial variability is clearly decreased after learning (as can be seen in fig. 4B). The observed increase in population-level variability in this set-up thus reflects a more constrained dynamical space and is a consequence of switching between highly active, specialized clusters. Thus, based on these observations from our model, we can expect that learning-induced modulations of population dynamics would result in an increase in population-level variability (as measured, for example by the CV). Having experimental access to a large enough population of responsive neurons would allow us to observe the formation of an increasingly restrictive dynamical space, whereby task-relevant variations would be imprinted in firing co-variation among strongly connected neuronal clusters (in line with Jazayeri & Afraz, 2017).

Our model constitutes an important step forward in the domain of unsupervised feature extraction, potentially leading to flexible learning algorithms, which can be used on large computational domains without requiring task-specific adjustments to the structure of the system. Because of its biological inspiration and plausibility, both in the learning rules and internal architecture, the model allows us to make predictions about the dynamic properties of internal representations and thus has the potential to lead to a better understanding of the principles underlying learning in the mammalian brain.

Capitalizing on these features, further work is required to clarify the extent of the model’s functional and biological relevance. For example, in this study we showed that the presence of clusters improved the performance of the network, but the number of clusters and their size relative to the whole network was fixed and chosen arbitrarily. It is reasonable to expect that the optimal parameter configuration would depend on the demands of the task. For example, data sets comprising larger input variability may benefit from a network architecture composed of a larger number of clusters. Future work should thus establish systematic relations between the number and size of internal network clusters, their functional impact relative to task demands as well as their effects on the observed dynamics. Furthermore, as we could use the same representational layer for different tasks, this suggests that this type of architecture may be suitable for multi-task learning. It remains to be seen how quickly the network can be re-trained on a different task, and whether such a re-mapping induces catastrophic forgetting of the first task, or permits a palimpsest of functionally relevant cluster mappings to be established.

## 5 ACKNOWLEDGMENTS

We acknowledge partial support by the German Federal Ministry of Education through our German-Japanese Computational Neuroscience Project (BMBF Grant 01GQ1343), the Helmholtz Alliance through the Initiative and Networking Fund of the Helmholtz Association and the Helmholtz Portfolio theme “Supercomputing and Modeling for the Human Brain”, and the European Union’s Horizon 2020 Framework Programme for Research and Innovation under Specific Grant Agreement Nos. 720270 and 785907 (Human Brain Project SGA1 and SGA2). We thank Dr. Vahid Rostami for sharing the PyNEST implementation of the clustered balanced random network and for all fruitful discussions.

## Footnotes

1 http://yann.lecun.com/exdb/mnist/

2 https://github.com/openai/gym/wiki/MountainCar-v0

3 https://fz-juelich.sciebo.de/s/iSAZ7be2prtCCXS

4 http://yann.lecun.com/exdb/mnist/

