## Supplementary Material for "Unsupervised learning and clustered connectivity enhance reinforcement learning in spiking neural networks"

### 1 Network description

| A | Model summary |
|---|---|
| **Populations** | Input, representation (excitatory and inhibitory) and output |
| **Connectivity** | Input to representation layer, Random recurrent connections in representation layer (with or without clusters), representation to output connections |
| **Neuron model** | Leaky integrate-and-fire, fixed voltage threshold, fixed absolute refractory time, exponential current-based synapses |
| **Synapse model** | Input: Hebbian with quadratic normalization term (Tetzlaff et al., 2013)<br>Output: Hebbian with quadratic normalization and neuromodulatory third factor<br>Reservoir: static |
| **Input** | Independent stochastic background input, rate-coded input signal (independent Poisson processes) |
| **Measurements** | Spiking activity, firing rates |

| B | Populations | |
|---|---|---|
| **Name** | **Elements** | **Size** |
| Background input | Independent Poisson processes | $n_\mathrm{b}$ (rate $F_\mathrm{b}$) |
| Input | Inhomogeneous Poisson processes | $n_\mathrm{inp}$ (rate $F_\mathrm{inp}$) |
| Reservoir exc | Integrate-and-fire neuron | $n_\mathrm{e}$ |
| Reservoir inh | Integrate-and-fire neuron | $n_\mathrm{i}$ |
| Output | Integrate-and-fire neuron | $n_\mathrm{out}$ |

| C | Connectivity | | |
|---|---|---|---|
| **Name** | **Source** | **Target** | **Pattern** |
| Input Projections | Input | Reservoir | all-to-all, initial weight $w_\mathrm{inp}$ |
| EE | Reservoir exc | Reservoir exc | Random pairwise Bernoulli, p = $p_\mathrm{ee}$, weight $w_\mathrm{ee}$ |
| EI | Reservoir exc | Reservoir inh | Random pairwise Bernoulli, p = $p_\mathrm{ei}$, weight $w_\mathrm{ei}$ |
| IE | Reservoir inh | Reservoir exc | Random pairwise Bernoulli, p = $p_\mathrm{ie}$, weight $w_\mathrm{ie}$ |
| II | Reservoir inh | Reservoir inh | Random pairwise Bernoulli, p = $p_\mathrm{ii}$, weight $w_\mathrm{ii}$ |
| Output Projections | Reservoir exc | Output | all-to-all, initial weight $w^\mu_\mathrm{out}$ $w^\sigma_\mathrm{out}$ |
| Background | Background input | Output | all-to-all, weight $w_\mathrm{b}$ |
| Output | Output | Output | all-to-all, weight $w_\mathrm{wta}$, delay $d_\mathrm{wta}$ |

| D | Neuron model |
|---|---|
| **Name** | `iaf_psc_exp` |
| **Subthreshold dynamics** | if $(t > t^* + \tau_\mathrm{ref})$ $\dot{V_m} = -V_m/\tau_m + (I_e + I_s)/C$ else $V_m(t) = V_\mathrm{reset}$<br>$\dot{I_s} = -I_s/\tau_s$ |
| **Spiking** | If $V(t-) < V_\mathrm{th}$ OR $V(t+) \geq V_\mathrm{th}$<br>1. set $t^* = t$<br>2. emit spike with time stamp $t^*$ |

| E | Plasticity | |
|---|---|---|
| **Source** | **Target** | **Equation** |
| Input | Reservoir | $\Delta w_{ij} = \mu \left( F_i F_j + \kappa (F^\mathrm{T} - F_j) w_{ij}^2 \right)$ |
| Reservoir | Output | $\Delta w_{ij} = \mu \left( (D - b_\mathrm{D}) F_i F_j + \kappa (F^\mathrm{T} - F_j) w_{ij}^2 \right)$ |

| F | Input | |
|---|---|---|
| **Source** | **Target** | **Description** |
| Poisson Generator | All reservoir and output | Independent background noise, rate $F_\mathrm{b}$, weight $w_\mathrm{b}$ |
| Inhomogeneous Poisson | Reservoir | Time-dependent rate (task-specific), initial weight $w_\mathrm{inp}$ |

| G | Measurements |
|---|---|
| Spiking activity (input, reservoir and output) | |

Table 1: Tabular description of network model after Nordlie et al., 2009.

### 1.1 Clustered balanced random network

| **A** | **Populations** | |
|---|---|---|
| **Name** | **Value** | **Description** |
| $n_{\mathrm{inp}}$ | task-dependent | Number of input neurons |
| $n_{\mathrm{e}}$ | 4000 | Number of excitatory neurons in representation layer |
| $n_{\mathrm{i}}$ | 1000 | Number of inhibitory neurons in representation layer |
| $n_{\mathrm{out}}$ | 3 | Number of output neurons |
| $n_{\mathrm{b}}$ | 1 | Number of background sources |
| **B** | **Connectivity** | |
| **Name** | **Value** | **Description** |
| $w_{\mathrm{inp}}$ | learned (initial value ) | Amplitude of excitatory input projections |
| $p_{\mathrm{ee}}^{\mathrm{intra}}$ | 0.2 | Connection probability for intra-cluster excitatory to excitatory connections |
| $w_{\mathrm{ee}}^{\mathrm{intra}}$ | 2.28 pA | Amplitude of intra-cluster excitatory to excitatory connections |
| $p_{\mathrm{ee}}^{\mathrm{extra}}$ | 0.2 | Connection probability for extra-cluster excitatory to excitatory connections |
| $w_{\mathrm{ee}}^{\mathrm{extra}}$ | 0.19 pA | Amplitude of extra-cluster excitatory to excitatory connections |
| $p_{\mathrm{ei}}^{\mathrm{intra}}$ | 0.5 | Connection probability for intra-cluster inhibitory to excitatory connections |
| $w_{\mathrm{ei}}^{\mathrm{intra}}$ | 1.33 pA | Amplitude of intra-cluster inhibitory to excitatory connections |
| $p_{\mathrm{ei}}^{\mathrm{extra}}$ | 0.5 | Connection probability for extra-cluster inhibitory to excitatory connections |
| $w_{\mathrm{ei}}^{\mathrm{extra}}$ | 0.19 pA | Amplitude of extra-cluster inhibitory to excitatory connections |
| $p_{\mathrm{ie}}^{\mathrm{intra}}$ | 0.5 | Connection probability for intra-cluster excitatory to inhibitory connections |
| $w_{\mathrm{ie}}^{\mathrm{intra}}$ | $-6.35$ pA | Amplitude of intra-cluster excitatory to inhibitory connections |
| $p_{\mathrm{ie}}^{\mathrm{extra}}$ | 0.5 | Connection probability for extra-cluster excitatory to inhibitory connections |
| $w_{\mathrm{ie}}^{\mathrm{extra}}$ | 0.91 pA | Amplitude of extra-cluster excitatory to inhibitory connections |
| $p_{\mathrm{ii}}^{\mathrm{intra}}$ | 0.5 | Connection probability for intra-cluster inhibitory to inhibitory connections |
| $w_{\mathrm{ii}}^{\mathrm{intra}}$ | $-9.24$ pA | Amplitude of intra-cluster inhibitory to inhibitory connections |
| $p_{\mathrm{ii}}^{\mathrm{extra}}$ | 0.5 | Connection probability for extra-cluster inhibitory to inhibitory connections |
| $w_{\mathrm{ii}}^{\mathrm{extra}}$ | 1.32 pA | Amplitude of extra-cluster inhibitory to inhibitory connections |
| $w_{\mathrm{out}}^{\mu}$ | 5.0 pA | Mean reservoir to output connection amplitude |
| $w_{\mathrm{out}}^{\sigma}$ | 1.44 | Standard deviation of reservoir to output connection amplitudes |
| $w_{\mathrm{wta}}$ | $-3000$ pA | Amplitude of output connections |
| $d_{\mathrm{wta}}$ | $[1, 3]$ ms | Delay of output connections |
| **D** | **Representation layer neurons** | |
| $C$ | 1 pF | Membrane capacitance |
| $V_{\mathrm{th}}$ | 20 mV | Fixed firing threshold |
| $\tau_{\mathrm{m}}$ (exc) | 20 ms | Membrane time constant (exc neurons) |
| $\tau_{\mathrm{m}}$ (inh) | 10 ms | Membrane time constant (inh neurons) |
| $\tau_{\mathrm{s}}$ | 2 ms | Synaptic time constant |
| $I_{\mathrm{e}}$ (exc) | 0.825 pA | Bias current (exc neurons) |
| $I_{\mathrm{e}}$ (inh) | 0.745 pA | Bias current (exc neurons) |
| $t_{\mathrm{ref}}$ | 5 ms | Absolute refractory period |
| **E** | **Output layer neurons** | |
| $C$ | 250 pF | Membrane capacitance |
| $V_{\mathrm{th}}$ | 15 mV | Fixed firing threshold |
| $\tau_{\mathrm{m}}$ | 20 ms | Membrane time constant |
| $\tau_{\mathrm{s}}$ | 2 ms | Synaptic time constant |
| $I_{\mathrm{e}}$ | 0 pA | Bias current |
| $t_{\mathrm{ref}}$ | 2 ms | Absolute refractory period |

Table 2: Model parameters for clustered networks.

### 1.2 Unclustered balanced random network

| A | | Populations |
|---|---|---|
| **Name** | **Value** | **Description** |
| $n_{\mathrm{inp}}$ | task-dependent | Number of input neurons |
| $n_{\mathrm{e}}$ | 4000 | Number of excitatory neurons in representation layer |
| $n_{\mathrm{i}}$ | 1000 | Number of inhibitory neurons in representation layer |
| $n_{\mathrm{out}}$ | 3 | Number of output neurons |
| $n_{\mathrm{b}}$ | 1 | Number of background sources |
| **B** | | **Connectivity** |
| **Name** | **Value** | **Description** |
| $w_{\mathrm{inp}}$ | task-dependent | Amplitude of excitatory input projections |
| $p_{\mathrm{ee}}$ | 0.2 | Connection probability for excitatory to excitatory connections |
| $w_{\mathrm{ee}}$ | 0.45 pA | Amplitude of excitatory to excitatory connections |
| $p_{\mathrm{ei}}$ | 0.5 | Connection probability for inhibitory to excitatory connections |
| $w_{\mathrm{ei}}$ | 0.33 pA | Amplitude of inhibitory to excitatory connections |
| $p_{\mathrm{ie}}$ | 0.5 | Connection probability for excitatory to inhibitory connections |
| $w_{\mathrm{ie}}$ | $-1.59$ pA | Amplitude of excitatory to inhibitory connections |
| $p_{\mathrm{ii}}$ | 0.5 | Connection probability for inhibitory to inhibitory connections |
| $w_{\mathrm{ii}}$ | $-2.31$ pA | Amplitude of inhibitory to inhibitory connections |
| $w_{\mathrm{out}}^{\mu}$ | 5.0 pA | Mean reservoir to output connection amplitude |
| $w_{\mathrm{out}}^{\sigma}$ | 1.44 | Standard deviation of reservoir to output connection amplitudes |
| $w_{\mathrm{wta}}$ | $-3000$ pA | Amplitude of output connections |
| $d_{\mathrm{wta}}$ | $[1, 3]$ ms | Delay of output connections |
| **D** | | **Representation layer neurons** |
| $C$ | 1 pF | Membrane capacitance |
| $V_{\mathrm{th}}$ | 20 mV | Fixed firing threshold |
| $\tau_{\mathrm{m}}$ (exc) | 20 ms | Membrane time constant (exc neurons) |
| $\tau_{\mathrm{m}}$ (inh) | 10 ms | Membrane time constant (inh neurons) |
| $\tau_{\mathrm{s}}$ | 2 ms | Synaptic time constant |
| $I_{\mathrm{e}}$ (exc) | 0.825 pA | Bias current (exc neurons) |
| $I_{\mathrm{e}}$ (inh) | 0.745 pA | Bias current (exc neurons) |
| $t_{\mathrm{ref}}$ | 5 ms | Absolute refractory period |
| **E** | | **Output layer neurons** |
| $C$ | 250 pF | Membrane capacitance |
| $V_{\mathrm{th}}$ | 15 mV | Fixed firing threshold |
| $\tau_{\mathrm{m}}$ | 20 ms | Membrane time constant |
| $\tau_{\mathrm{s}}$ | 2 ms | Synaptic time constant |
| $I_{\mathrm{e}}$ | 0 pA | Bias current |
| $t_{\mathrm{ref}}$ | 2 ms | Absolute refractory period |

Table 3: Model parameters for unclustered networks.

### 2 Task description

| XOR | | |
|---|---|---|
| **Name** | **Value** | **Description** |
| $F_{\text{inp}}$ | $[10, 1000]$ Hz | Range of input firing rates |
| $w_{\text{inp}}$ | 50 pA | Initial amplitude of input projections |
| **Input Plasticity** | | |
| $\mu$ | 0.000001 | Global learning rate for input synapses |
| $\kappa$ | 1.0 | Ratio of synaptic scaling to Hebbian learning |
| $F^{\text{T}}$ | 0.15 | Homeostatic set point (target firing rate) |

| MNIST | | |
|---|---|---|
| **Name** | **Value** | **Description** |
| $F_{\text{b}}$ | 95 kHz | Background firing rate |
| $w_{\text{b}}$ | 2 pA | Amplitude of background input synapses |
| **Input** | | |
| $F_{\text{inp}}$ | $[0, 20]$ Hz | Range of input firing rates |
| $w_{\text{inp}}$ | 3 pA | Initial amplitude of input projections |
| **Input Plasticity** | | |
| $\mu$ | 0.0001 | Global learning rate |
| $\kappa$ | 0.2 | Ratio of synaptic scaling to Hebbian learning |
| $F^{\text{T}}$ | 0.1 | Homeostatic set point (target firing rate) |
| **Output Plasticity** | | |
| $\mu$ | 0.00001 | Global learning rate |
| $\kappa$ | 0.0 | Ratio of synaptic scaling to Hebbian learning |
| $F^{\text{T}}$ | 0.0 | Homeostatic set point (target firing rate) |
| $b_{\text{D}}$ | 9.7 | Baseline dopaminergic concentration |

| Mountain Car | | |
|---|---|---|
| **Name** | **Value** | **Description** |
| $F_{\text{b}}$ | 95 kHz | Background firing rate |
| $w_{\text{b}}$ | 2 pA | Amplitude of background input synapses |
| **Input** | | |
| $F_{\text{inp}}$ | $[0, 35]$ Hz | Range of input firing rates |
| $w_{\text{inp}}$ | 2 pA | Initial amplitude of input projections |
| **Input Plasticity** | | |
| $\mu$ | 0.0001 | Global learning rate |
| $\kappa$ | 0.12 | Ratio of synaptic scaling to Hebbian learning |
| $F^{\text{T}}$ | 0.1 | Homeostatic set point (target firing rate) |
| **Output Plasticity (actor)** | | |
| $\mu$ | 0.00001 | Global learning rate |
| $\kappa$ | 0.0 | Ratio of synaptic scaling to Hebbian learning |
| $F^{\text{T}}$ | 0.0 | Homeostatic set point (target firing rate) |
| $b_{\text{D}}$ | 9.7 | Baseline dopaminergic concentration |
| **Output Plasticity (critic)** | | |
| $\mu$ | 0.00001 | Global learning rate |
| $\kappa$ | 0.0 | Ratio of synaptic scaling to Hebbian learning |
| $F^{\text{T}}$ | 0.0 | Homeostatic set point (target firing rate) |
| $b_{\text{D}}$ | 9.7 | Baseline dopaminergic concentration |
